# SARS-CoV-2 selectively mimics a cleavable peptide of human ENaC in a strategic hijack of host proteolytic machinery

**DOI:** 10.1101/2020.04.29.069476

**Authors:** Praveen Anand, Arjun Puranik, Murali Aravamudan, AJ Venkatakrishnan, Venky Soundararajan

## Abstract

Molecular mimicry of host proteins is an evolutionary strategy adopted by viruses to evade immune surveillance and exploit host cell systems. We report that SARS-CoV-2 has evolved a unique S1/S2 cleavage site (RRARSVAS), absent in any previous coronavirus sequenced, that results in mimicry of an identical FURIN-cleavable peptide on the human epithelial sodium channel α-subunit (ENaC-α). Genetic truncation at this ENaC-α cleavage site causes aldosterone dysregulation in patients, highlighting the functional importance of the mimicked SARS-CoV-2 peptide. Single cell RNA-seq from 65 studies shows significant overlap between the expression of ENaC-α and ACE2, the putative receptor for the virus, in cell types linked to the cardiovascular-renal-pulmonary pathophysiology of COVID-19. Triangulating this cellular fingerprint with amino acid cleavage signatures of 178 human proteases shows the potential for tissue-specific proteolytic degeneracy wired into the SARS-CoV-2 lifecycle. We extrapolate that the evolution of SARS-CoV-2 into a global coronavirus pandemic may be in part due to its targeted mimicry of human ENaC and hijack of the associated host proteolytic network.

---

The surface of SARS-CoV-2 virions is coated with the spike (S) glycoprotein, whose proteolysis is key to the infection lifecycle. After the initial interaction of the S-protein with the ACE2 receptor^1^, host cell entry is mediated by two key proteolytic steps. The S1 subunit of the S-protein engages ACE2, and viral entry into the host cell is facilitated by proteases that catalyze S1/S2 cleavage^2,3^ at Arginine-667/Serine-668 (**Figure 1a**). This is followed by S2’ site cleavage that is required for fusion of viral-host cell membranes^1^.

**Figure 1.**
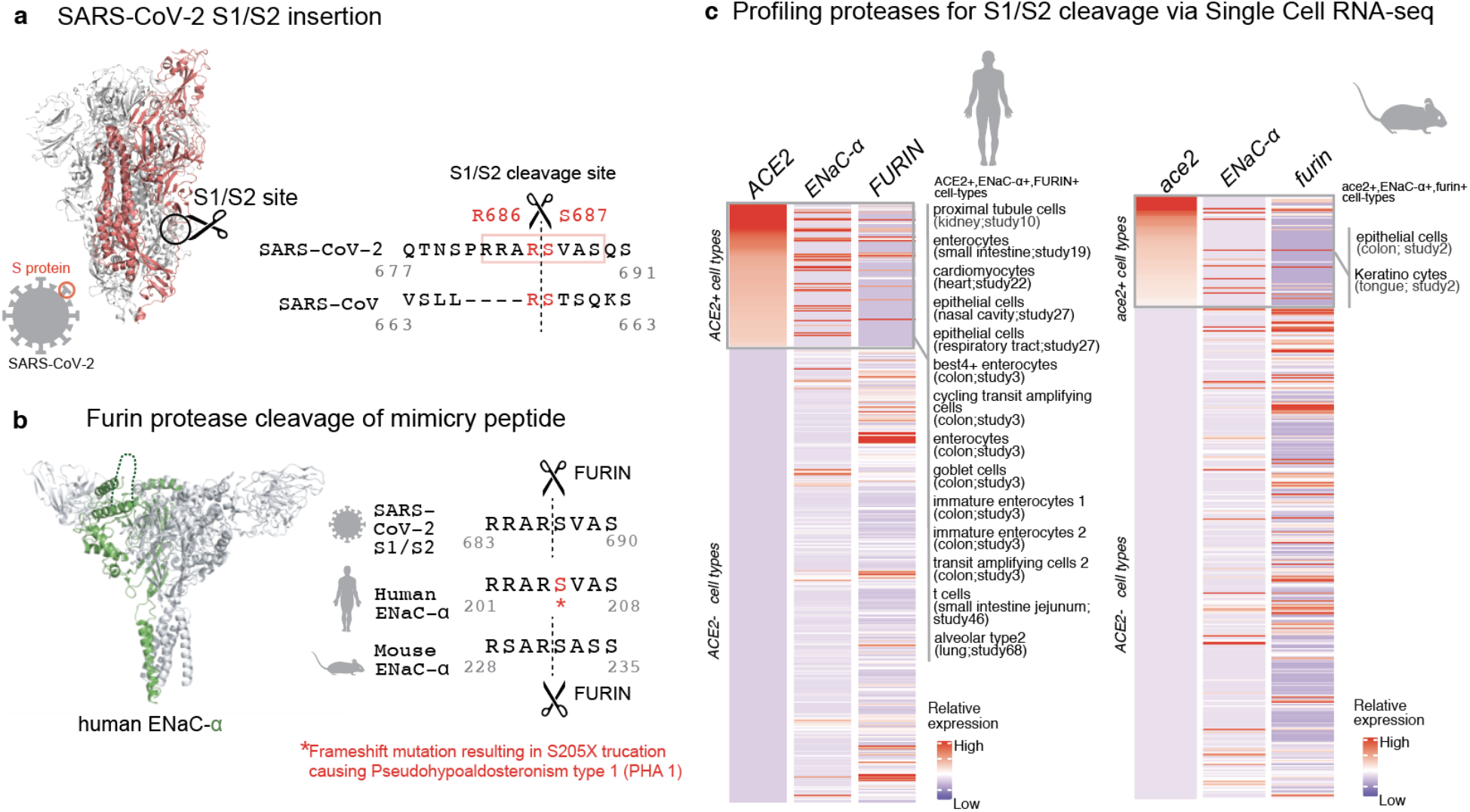
Targeted molecular mimicry by SARS-CoV-2 of human ENaC-ɑ and profiling ACE2-FURIN-ENaC-ɑ co-expression. **(a)** The cartoon representation of the S-protein homotrimer from SARS-CoV-2 is shown (PDB ID: 6VSB). One of the monomers in highlighted red. The pairwise alignment of the S1/S2 cleavage site required for the activation of SARS and SARS-CoV-2 is depicted. The 4 amino acid insertion evolved by SARS-CoV-2, along with the abutting cleavage site is shown in a box. **(b)** The cartoon representation of human ENaC protein is depicted (PDB ID: 6BQN; chain in green), highlighting the ENaC-ɑ chain in green. The alignment on the right captures FURIN cleavage at the S1/S2 site of SARS-CoV-2, along with its striking molecular mimicry of the identical peptide from human ENaC-ɑ protein (circled in the cartoon rendering of human ENaC). The alignment also shows the equivalent 8-mer peptide of mouse ENaC-ɑ that is also known to be cleaved by FURIN. **(c)** The single cell transcriptomic co-expression of ACE2, ENaC-ɑ, and FURIN is summarized. The heatmap depicts the mean relative expression of each gene across the identified cell populations. The human and mouse single cell RNA-seq are visualized independently. The cell types are ranked based on decreasing expression of ACE2. The box highlights the ACE2 positive cell types in mouse and human samples.

We hypothesized that the virus may mimic host substrates in order to achieve proteolysis. Comparing SARS-CoV-2 with SARS-CoV shows that the former has acquired a distinctive sequence insertion at the S1/S2 site (**Figure 1a**). The resulting tribasic 8-mer peptide (RRARSVAS) on the SARS-CoV-2 S1/S2 site is conserved among 10,956 of 10,967 circulating strains deposited at GISAID^4^, as of April 28, 2020 (*Table S1*). This peptide is also absent in over 13,000 non-COVID-19 coronavirus S-proteins from the VIPR database^5^. Strikingly, examining over 10 million peptides (8-mers) of 20,350 canonical human proteins from UniProtKB shows that the peptide of interest (RRARSVAS) is present exclusively in human ENaC-ɑ, also known as SCNN1A (p-value = 4E-4) (*Supplementary Methods*). The location of this SARS-CoV-2 mimicked peptide in the ENaC-ɑ structure is in the extracellular domain (**Figure 1b**).

ENaC regulates Na+ and water homeostasis and its expression levels are controlled by aldosterone and the associated Renin-Angiotensin-Aldosterone System (RAAS)^6^. Similar to SARS-CoV2, ENaC-ɑ needs to be proteolytically activated for its function^7^. FURIN cleaves the equivalent peptide on mouse ENaC-ɑ between the Arginine and Serine residues in the 4^th^ and 5^th^ positions respectively (RSAR|SASS)^8,9^, akin to the recent report establishing FURIN cleavage at the S1/S2 site of SARS-CoV-2 (**Figure 1b**)^1^. Furthermore, a frameshift mutation leading to a premature stop codon in Serine-205 at the 5^th^ position of the ENaC-ɑ mimicked peptide (RRAR|<u>S</u>VAS) is known to cause the monogenic disease Pseudohypoaldosteronism type 1 (PHA1)^10^. This emphasizes the functional salience of the 8-mer peptide being mimicked by SARS-CoV-2.

Mimicry of human ENaC-ɑ by the S1/S2 site raises the specter that SARS-CoV-2 may be hijacking the protease network of ENaC-ɑ for viral activation. We asked whether there is an overlap between putative SARS-CoV-2 infecting cells and ENaC-ɑ expressing cells. Systematic single cell expression profiling of the ACE2 receptor and ENaC-ɑ was performed across human and mouse samples comprising ~1.3 million cells (*Supplementary Methods*, **Figure 1c**)^11^. Interestingly, ENaC-ɑ is expressed in the nasal epithelial cells, type II alveolar cells of the lung, tongue keratinocytes, and colon enterocytes (**Figure 1c** and *Figures S1-S6*), which are all implicated in COVID-19 pathophysiology^11,12^. Further, ACE2 and ENaC-ɑ are known to be expressed generally in the apical membranes of polarized epithelial cells^13,14^. The overlap of the cell-types expressing ACE2 and ENaC-ɑ, and similar spatial distributions at the apical surfaces, suggest that SARS-CoV-2 may be leveraging the protease network responsible for ENaC cleavage.

Beyond FURIN that cleaves the S1/S2 site^1^, we were intrigued by the possibility of other host proteases also being exploited by SARS-CoV-2. We created a 160-dimensional vector space (20 amino acids x 8 positions on the peptide) for assessment of cleavage similarities between the 178 human proteases with biochemical validation from the MEROPS database (*Supplementary Methods*; 0 < protease similarity metric < 1)^15^. This shows that FURIN (PCSK3) has overall proteolytic similarity to select PCSK family members, specifically PCSK5 (0.99), PCSK7 (0.99), PCSK6 (0.99), PCSK4 (0.98), and PCSK2 (0.94) (*Table S2*). It is also known that the protease PLG cleaves the ɣ-subunit of ENaC (ENaC-ɣ)^16^.

In order to extrapolate the tissue tropism of SARS-CoV-2 from the lens of the host proteolytic network, we assessed the co-expression of these proteases concomitant with the viral receptor ACE2 and ENaC-ɑ (**Figure 2**). This analysis shows that FURIN is expressed with ACE2 and ENaC-ɑ in the colon (immature enterocytes, transit amplifying cells) and pancreas (ductal cells, acinar cells) of human tissues, as well as tongue (keratinocytes) of mouse tissues. PCSK5 and PCSK7 are broadly expressed across multiple cell types with ACE2 and ENaC-ɑ, making it a plausible broad-spectrum protease that may cleave the S1/S2 site. In humans, concomitant with ACE2 and ENaC-ɑ, PCSK6 appears to be expressed in cells from the intestines, pancreas, and lungs, whereas PCSK2 is noted to be co-expressed in the respiratory tract and the pancreas (**Figure 2**). It is worth noting that the extracellular proteases need not necessarily be expressed in the same cells as ACE2 and ENaC-ɑ. Among the PCSK family members with the potential to cleave the mimicked 8-mer peptide, it is intriguing that the same tissue can house multiple proteases and also that multiple tissues do share the same set of proteases.

**Figure 2.**
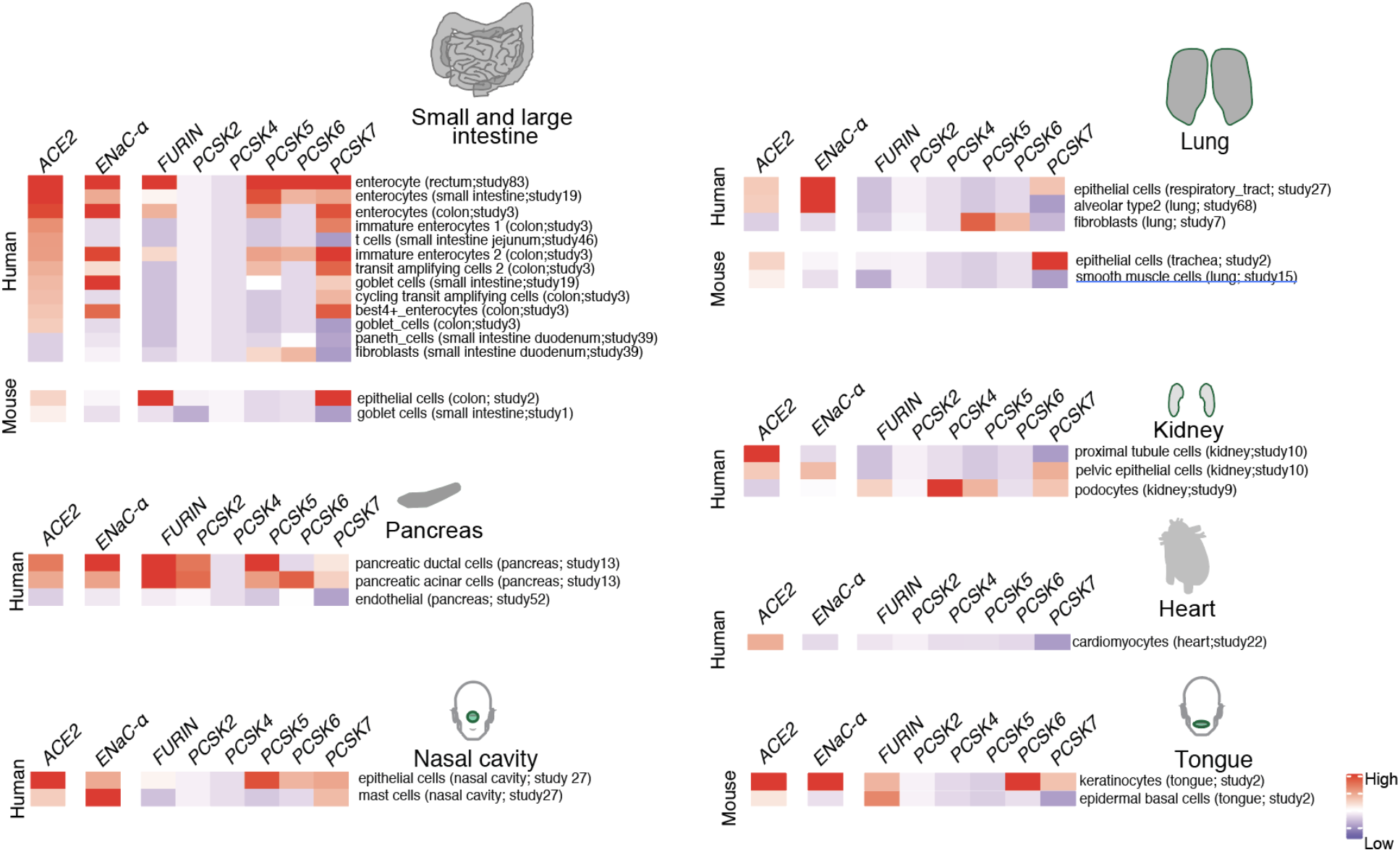
Expression profiling of identified proteases. The heatmap depicts the relative expression of ACE2 and ENaC-ɑ along with all the proteases that can potentially cleave the S1/S2 site. The relative expression levels are denoted on a scale of blue (low) to red (high). The rows denote proteases and columns denote cell-types.

Our findings emphasize that redundancy may be wired into the mechanisms of host proteolytic activation of SARS-CoV-2, and call for some caution in the ongoing development of selective human protease inhibitors as COVID-19 therapeutics. The mimicry of a cleavable host peptide central to cardiovascular, renal, and pulmonary function provides a new perspective to the evolution of SARS-CoV-2 as the first coronavirus pandemic.

## Acknowledgements

The authors thank Patrick Lenehan, David Zemmour, Travis Hughes, Tyler Wagner, and Mathai Mammen for their careful review and feedback. The authors are also grateful to Ramakrishna Chilaka for the software visualization tools, and Dhruti Patwardhan, Saranya Marimuthu, Jaya Jain, Dariusz Murakowski, and Enrique Garcia-Rivera for their assistance with databases.

