## Supplementary Material for "SARS-CoV-2 selectively mimics a cleavable peptide of human ENaC in a strategic hijack of host proteolytic machinery"

#### Supplementary Methods

##### Alignment of coronavirus spike proteins

The complete S-protein sequence for SARS-CoV (Uniprot ID: P59594) and SARS-CoV-2 was obtained from uniprot (ftp://ftp.uniprot.org/pub/databases/uniprot/pre_release/)[16]. Sequence alignments using Clustal-W, and comparison of SARS-CoV-2 versus other coronavirus strains were performed using JalView[17].

##### 8-mer analysis of human proteome

The number of 8-mers in Uniprot 20,350 reference sequences are 10,257,893 (10.26M). The previously identified SARS-CoV-2 8-mer 'RRARSVAS' was in fact found in a Uniprot reference sequence (p-value ≈ $10.26M/20^8 = 4E-4$; chance of finding that particular 8-mer anywhere in the reference sequences).

##### Scoring the cleavage sites

The biochemical specificities are inferred from substrates of the proteases that have been determined using the evidence from (i) N-terminal sequencing, (ii) mass spectroscopy, (iii) mutational studies, (iv) consensus analysis or (v) liquid chromatography. The protein residues from the protease substrates spanning the scissile bond (±4 residues) is considered to define sequence specificity of the motif. The motif length thus spans across 8 residues (or positions), and frequency of all the 20 amino acids at each position is calculated from the substrates identified for each of the proteases to get a position frequency matrix (PFM, with a dimension of 8 x 20). This PFM was converted to a probability matrix by normalizing to the frequency distribution of all the 20 amino acids per position. The position frequency matrices were downloaded from the MEROPS database[15]. In total, probabilities matrices for 178 proteases were constructed. The number of distinct cleavages for each protease ranges from as few as 10 to over 3000, based on which probability matrices for the cleavage of the 8-mer polypeptide was estimated (**Table S2**). The probability matrix is then used to scan the peptide sequence ('RRARSVAS') using a window frame equal to the length of the motif. A log-likelihood ratio score is calculated for each sequence position in the scanned window with respect to background distribution of particular amino acid in the human proteome and summed up across all the positions to get a motif score. A random set of sequences are generated from the given background of amino acid distribution in the human proteome and used as a null model to evaluate the statistical significance of the scanned window. FIMO tool was used to obtain the scores and the corresponding statistical significance of the PWM match[18]. These were then used to estimate cleavage propensity by assuming background distribution of amino-acid frequencies from human proteome[19] using MEME[20] and FIMO[18]. A stringent cut-off of 1E-04 was used to identify the potential cleavage sites on S proteins of both SARS-CoV and SARS-CoV-2 at S1/S2 site. The hits obtained were further manually filtered to ensure that residue at the immediate vicinity of cleavage site is conserved.

### Supplementary Material

#### Calculating the cosine similarity metric for protease cleavage site

The position frequency matrix (PFM) of the individual proteases obtained from the MEROPS database was converted to a probability weight matrix (PWM) (normalized and scaled). Out of 178 proteases, there were 146 proteases that had specificity information available on the 8 mer peptide spanning the cleavage site (±4). The 20 (amino acids) x 8 (position) matrix defined for each of the proteases were flattened into a single vector with 160 elements. We performed a cosine similarity calculation between all pairs (X,Y) of protease specificity vector. The similarity was derived as the normalized dot product of X and Y : $K(X, Y) = \langle X, Y \rangle / (||X||*||Y||)$.

#### Overlap of cell types expressing ENaC-α, ACE2 and proteases from scRNA-seq datasets

We performed a systematic expression profiling of the ACE2 and ENaC-α across 65 published human and mouse single-cell studies comprising ~1.3 million cells using nferX Single Cell platform (**Table S3**, https://academia.nferx.com/)[6]. The ACE2 expression could be detected in 67 studies (59 studies of human samples and 8 studies of mouse samples) spanning across ~50 tissues, over 450 cell-types and ~1.05 million cells. In order to call a given cell-type to be positive for both ACE2 and a protease we applied a cutoff of 1% of the cells in the total cell-type cluster population to have a non-zero count associated with both ACE2 and the respective protease. The mean expression of the proteases, ENaC-α and ACE2 was derived for individual cell population within each of the studies. The cell-type information was obtained from the author annotations provided for each of the studies. The analysis was performed separately on the mouse and human datasets. For each protease, the mean expression of in a given cell-population (mean log[cp10k +1] counts) was Z-score normalized (to ensure the sd=1 and mean ~0 for all the genes) to obtain relative expression profiles across all the samples. The same normalization was applied to ACE2 and ENaC-α and both human and mouse datasets were analyzed independently by generating heatmaps. The cell types having zero-expression values of ACE2 were also included as negative control to probe the expression of various proteases.

We performed an analysis to identify the cell types with significant overlap of ACE2 and ENaC-α expression. To this end, we shortlisted cell types in which ENaC-α is expressed in a significantly higher proportion of ACE2-expressing cells than in the overall population of cells of that sub-type. We computed the ratios of these proportions, and used a corresponding Fisher exact test to compute significance.

### Supplementary Material

**Table S1.** SARS-CoV-2 variants in the RRARSVAS 8-mer peptide from 10,987 spike (S) protein sequences of the GISAID database. The specific variations are highlighted in **Red**.

| Variation in the mimicked 8-mer of interest (RRARSVAS) | Number of occurrences in the SARS-CoV-2 S-protein sequences | Strain Information (GISAID) |
|---|---|---|
| **RRARSVAS** | **10,976** | - |
| RPARSVAS | 1 | HCOV-19/NETHERLANDS/ZUIDHOLLAND_37/2020\|EPI_ISL_422909\|2020-03-17 |
| QRARSVAS | 1 | HCOV-19/HANGZHOU/ZJU-01/2020\|EPI_ISL_415709\|2020-01-25 |
| RQARSVAS | 1 | HCOV-19/ENGLAND/CAMB-73800/2020\|EPI_ISL_425243\|2020-04-01 |
| RRARGVAS | 1 | HCOV-19/RUSSIA/KRASNODAR-63401/2020\|EPI_ISL_428867\|2020-03-11 |
| RRARSVVS | 2 | HCOV-19/ENGLAND/20104035803/2020\|EPI_ISL_417238\|2020-03-0<br>HCOV-19/WALES/PHWC-2658D/2020\|EPI_ISL_422346\|2020-03-26 |
| RRARSVAI | 3 | HCOV-19/ENGLAND/20140007302/2020\|EPI_ISL_421925\|2020-03-28<br>HCOV-19/ENGLAND/20140005304/2020\|EPI_ISL_423380\|2020-03-29<br>'HCOV-19/FRANCE/ARA12265/2020\|EPI_ISL_419186\|2020-03-22 |
| RRVRSVAS | 2 | HCOV-19/BRAZIL/RJ-872/2020\|EPI_ISL_427304\|2020-03-26<br>HCOV-19/SPAIN/VALENCIA98/2020\|EPI_ISL_425222\|2020-03-17 |
| **[RQ][RQP][AV][R][SG][V][AV][IS]** | **10,987** | |

### Supplementary Material

**Table S2.** Protease cleavage propensities for FURIN and the other proteases identified as similar from the vector space analysis conducted. Similarity (FURIN) ranges from 0 to 1. Highlighted green are amino acids occurring in greater than 10% of the cleaved substrates at that position (compiled from MEROPS).

| Protease | Cleavage substrates | Similarity (FURIN) | P4 | P3 | P2 | P1 | P1' | P2' | P3' | P4' |
|---|---|---|---|---|---|---|---|---|---|---|
| MIMICKED PEPTIDE | | | R | R | A | R | S | V | A | S |
| **FURIN** | 208 | 1.00 | R(158)<br>I(8)<br>K(7)<br>F(7)<br>Others(26) | K(34)<br>S(27)<br>R(26)<br>T(18)<br>Others(97) | K(88)<br>R(68)<br>P(9)<br>A(8)<br>Others(34) | R(203)<br>K(4)<br>L(1) | S(57)<br>A(23)<br>D(22)<br>E(20)<br>Others(86) | V(46)<br>A(34)<br>L(31)<br>I(15)<br>Others(76) | S(30)<br>G(21)<br>D(18)<br>E(16)<br>Others(113) | S(19)<br>G(19)<br>A(17)<br>E(16)<br>Others(129) |
| **PCSK5** | 129 | 0.992 | R(97)<br>K(8)<br>I(6)<br>V(4)<br>Others(12) | K(23)<br>S(16)<br>R(13)<br>Q(10)<br>Others(62) | K(59)<br>R(41)<br>P(8)<br>S(4)<br>Others(12) | R(125)<br>K(4) | S(37)<br>A(11)<br>D(11)<br>F(9)<br>Others(58) | V(25)<br>A(24)<br>L(22)<br>I(12)<br>Others(43) | S(15)<br>G(15)<br>D(15)<br>E(13)<br>Others(62) | E(15)<br>L(13)<br>P(12)<br>G(11)<br>Others(71) |
| **PCSK4** | 103 | 0.99 | R(77)<br>K(8)<br>V(4)<br>I(2)<br>Others(7) | K(18)<br>R(12)<br>S(11)<br>Q(10)<br>Others(45) | K(49)<br>R(32)<br>P(6)<br>A(3)<br>Others(7) | R(100)<br>K(3) | S(31)<br>E(9)<br>D(9)<br>A(9)<br>Others(37) | V(25)<br>A(19)<br>L(15)<br>T(10)<br>Others(26) | S(13)<br>G(12)<br>D(12)<br>E(11)<br>Others(47) | E(13)<br>P(11)<br>L(11)<br>S(8)<br>Others(52) |
| **PCSK6** | 105 | 0.99 | R(85)<br>K(7)<br>V(4)<br>I(2)<br>Others(7) | K(19)<br>S(12)<br>R(12)<br>Q(10)<br>Others(45) | K(53)<br>S(36)<br>R(6)<br>Q(3)<br>Others(7) | R(102)<br>K(3) | S(33)<br>A(10)<br>E(9)<br>D(9)<br>Others(41) | V(29)<br>A(20)<br>L(15)<br>T(10)<br>Others(28) | G(15)<br>S(14)<br>D(13)<br>E(11)<br>Others(49) | E(13)<br>L(12)<br>P(11)<br>S(10)<br>Others(46) |
| **PCSK7** | 117 | 0.989 | R(85)<br>K(9)<br>I(5)<br>V(4)<br>Others(8) | K(23)<br>S(13)<br>R(12)<br>Q(11)<br>Others(50) | K(54)<br>R(38)<br>P(7)<br>A(3)<br>Others(8) | R(112)<br>K(4) | S(34)<br>E(11)<br>D(10)<br>A(10)<br>Others(44) | V(25)<br>L(22)<br>A(20)<br>T(11)<br>Others(31) | D(14)<br>S(13)<br>G(13)<br>E(13)<br>Others(52) | E(15)<br>P(11)<br>L(11)<br>A(10)<br>Others(60) |
| **PCSK2** | 205 | 0.941 | R(86)<br>K(13)<br>V(11)<br>I(11)<br>Others(8) | Q(27)<br>S(22)<br>K(20)<br>E(19)<br>Others(109) | K(123)<br>R(44)<br>P(9)<br>A(6)<br>Others(9) | R(192)<br>K(11)<br>S(1)<br>F(1) | S(43)<br>Y(25)<br>A(20)<br>G(15)<br>Others(93) | V(27)<br>G(23)<br>L(22)<br>A(22)<br>Others(102) | G(31)<br>E(27)<br>S(17)<br>Q(17)<br>Others(103) | E(31)<br>D(27)<br>F(17)<br>S(17)<br>Others(117) |
| **PLG** | 126 | | P(18)<br>A(16)<br>R(13)<br>S(8)<br>Others(52) | R(17)<br>S(12)<br>Q(11)<br>G(10)<br>Others(70) | L(15)<br>S(13)<br>P(12)<br>A(11)<br>Others(72) | R(65)<br>K(57)<br>Others(3) | S(23)<br>A(20)<br>G(11)<br>R(10)<br>Others(51) | R(13)<br>V(12)<br>S(12)<br>K(8)<br>Others(70) | S(13)<br>P(11)<br>A(9)<br>Q(8)<br>Others(74) | G(12)<br>P(11)<br>L(11)<br>A(9)<br>Others(72) |

### Supplementary Material

#### Supplementary Figures

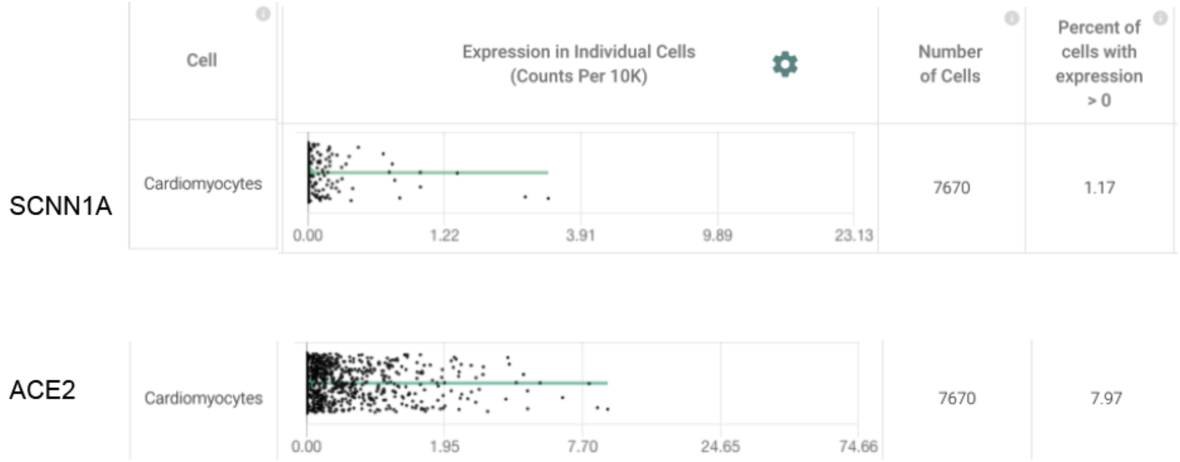


**Figure S1.** Cardiomyocytes express ENaC-α (SCNN1A) and ACE2 (Primary data processed from Pubmed ID:31915373 and hosted on [https://academia.nferx.com/](https://academia.nferx.com/))

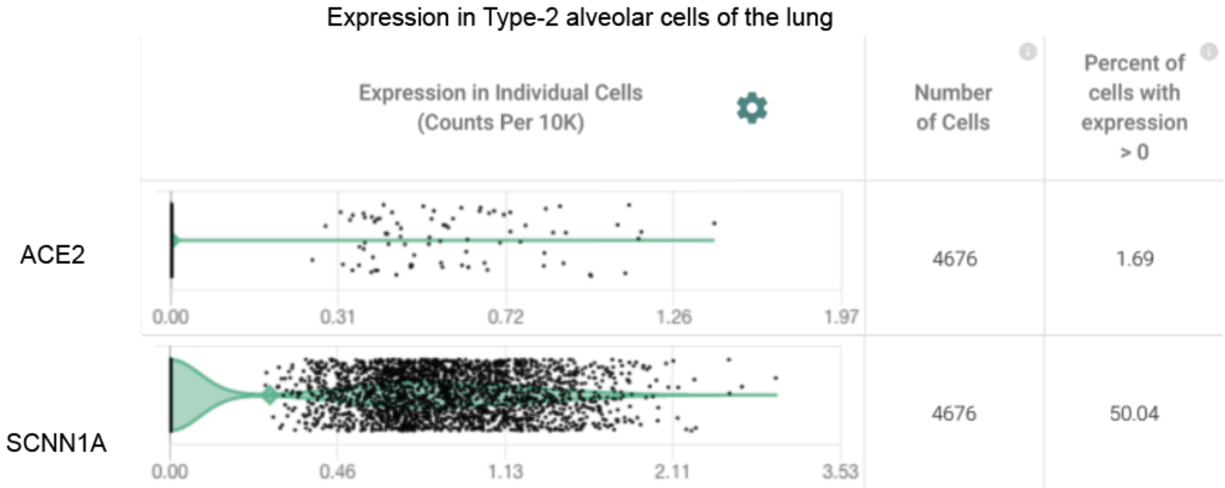


**Figure S2.** Type-II Alveolar Cells of the lungs express ENaC-α (SCNN1A) and ACE2 (Primary data processed from Pubmed ID: 31892341 and hosted on [https://academia.nferx.com/](https://academia.nferx.com/))

### Supplementary Material

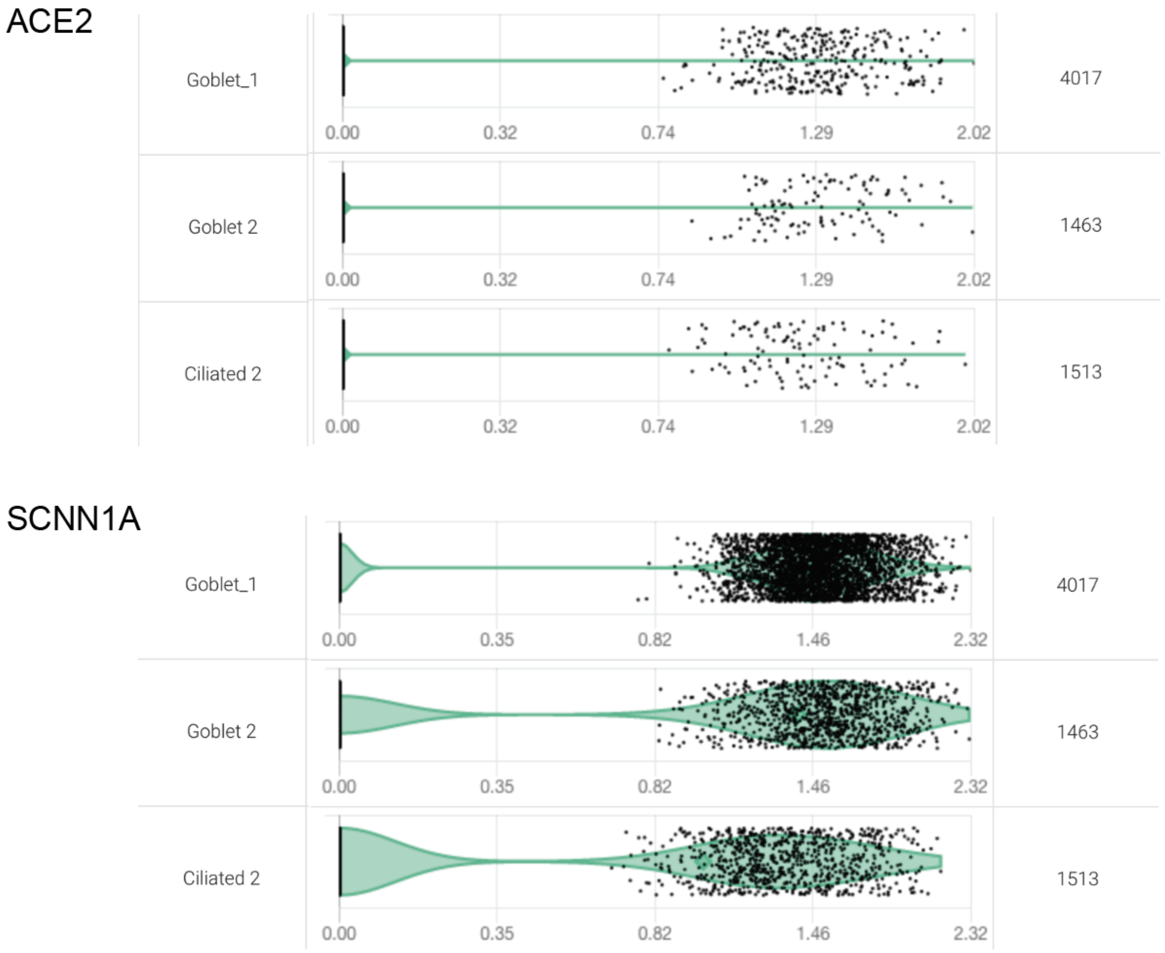


**Figure S3.** Goblet cells and Ciliated cells of the nasal epithelial layer express SCNN1A (ENaC-α) and ACE2 (Primary data processed from Pubmed ID: 32327758 and hosted on https://academia.nferx.com/)

### Supplementary Material

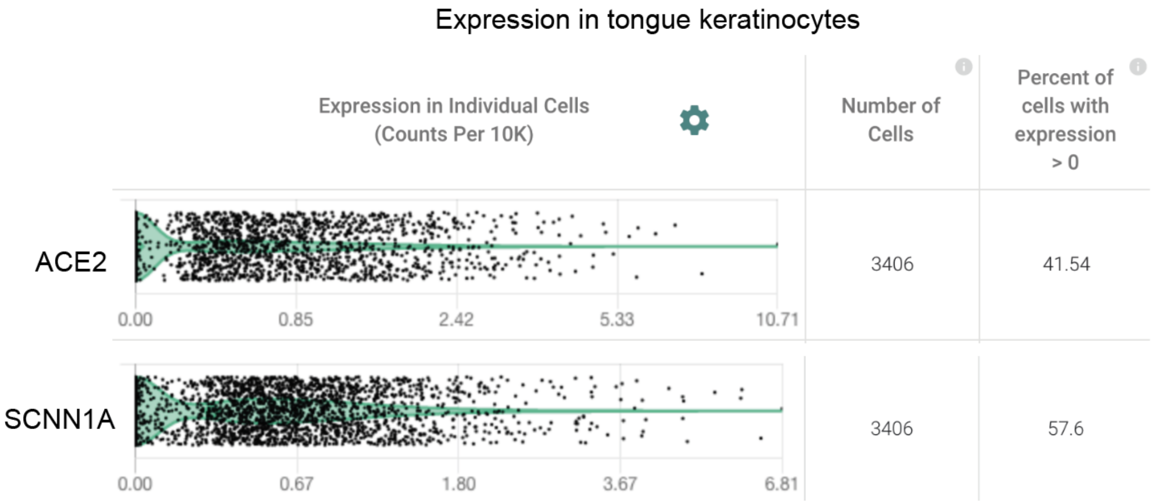


**Figure S4.** Tongue keratinocytes express SCNN1A (ENaC-α) and ACE2 (Primary data processed from Pubmed ID:30283141 and hosted on https://academia.nferx.com/)

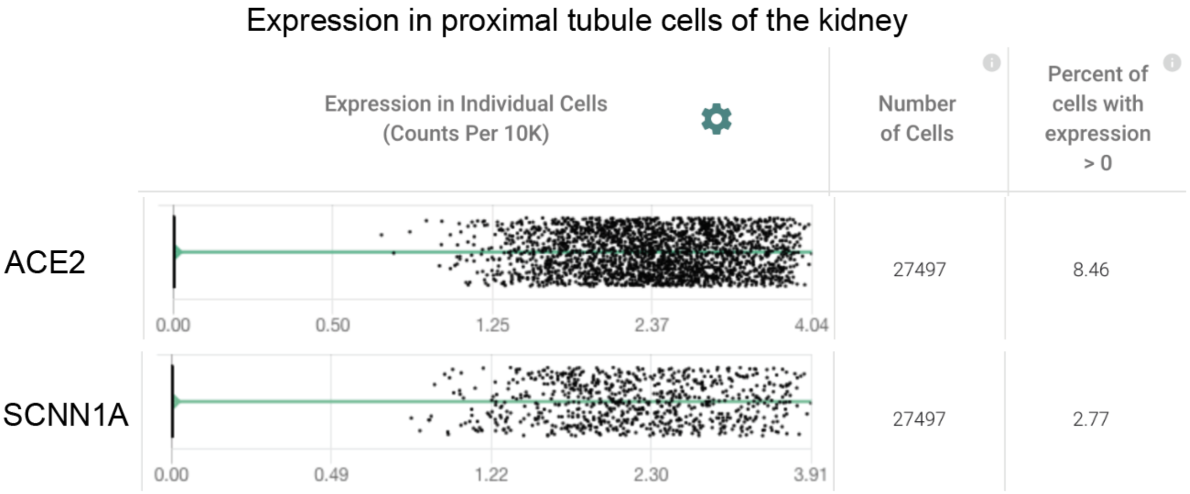


**Figure S5.** Kidney proximal tubule cells express SCNN1A (ENaC-α) and ACE2 (Primary data processed from Pubmed ID: 31604275 and hosted on https://academia.nferx.com/)

### Supplementary Material

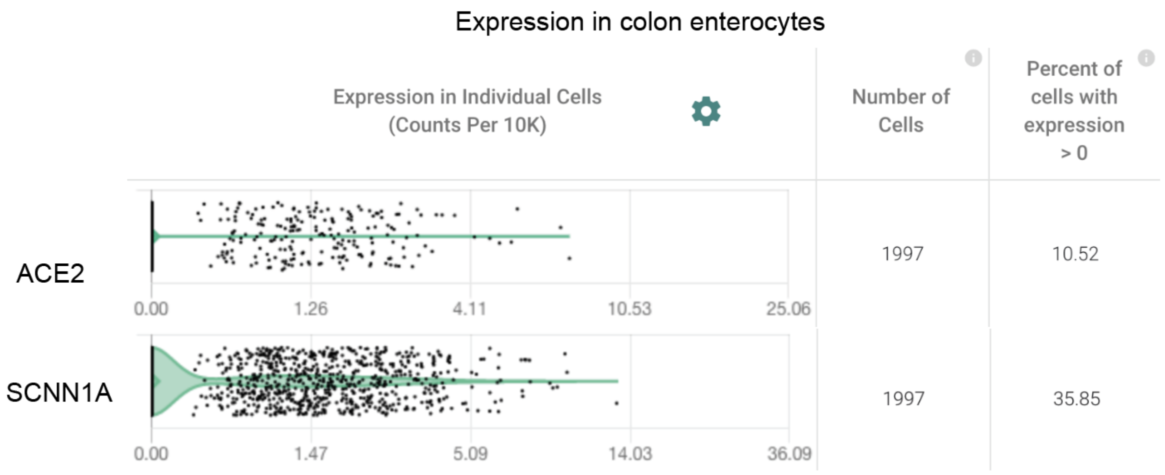


**Figure S6.** Colon enterocytes express SCNN1A (ENaC-α) and ACE2 (Primary data processed from Pubmed ID:31348891 and hosted on https://academia.nferx.com/)

### Supplementary Material

**Table S3. List of single-cell studies analyzed and incorporated into the nferX resource (https://academia.nferx.com/)**

| Study ID | Organism | Study Title | Pubmed ID (PMID) |
|---|---|---|---|
| study1 | Mus musculus | A single-cell survey of the small intestinal epithelium | PMID: 29144463 |
| study2 | Mus musculus | Single-cell transcriptomics of 20 mouse organs creates a Tabula Muris. | PMID:30283141 |
| study3 | Homo sapiens | Intra- and Inter-cellular Rewiring of the Human Colon during Ulcerative Colitis | PMID:31348891 |
| study4 | Homo sapiens | Immune Cell Atlas: Blood Mononuclear Cells (2 donors, 2 sites) | https://singlecell.broadinstitute.org/single_cell/study/SCP345/ica-blood-mononuclear-cells-2-donors-2-sites |
| study5 | Homo sapiens | Spleen - Ischaemic sensitivity of human tissue by single cell RNA seq | https://data.humancellatlas.org/explore/projects/c4077b3c-5c98-4d26-a614-246d12c2e5d7 |
| study6 | Homo sapiens | Esophagus - Ischaemic sensitivity of human tissue by single cell RNA seq | https://data.humancellatlas.org/explore/projects/c4077b3c-5c98-4d26-a614-246d12c2e5d7 |
| study7 | Homo sapiens | A cellular census of human lungs identifies novel cell states in health and in asthma. | PMID: 31209336 |
| study8 | Mus musculus | A revised airway epithelial hierarchy includes CFTR-expressing ionocytes | PMID: 30069044 |
| study9 | Homo sapiens | Fetal Kidney - Spatiotemporal immune zonation of the human kidney | PMID: 31604275 |
| study10 | Homo sapiens | Mature Kidney - Spatiotemporal immune zonation of the human kidney | PMID: 31604275 |
| study11 | Homo sapiens | Identification of grade and origin specific cell populations in serous epithelial ovarian cancer by single cell RNA-seq | PMID: 30383866 |
| study12 | Homo sapiens | A human liver cell atlas reveals heterogeneity and epithelial progenitors. | PMID:31292543 |
| study13 | Homo sapiens | Human Pancreas scRNA-seq (Integration of 3 Datasets) | PMID:27345837,PMID:27667667,PMID:27693023 |
| study14 | Homo sapiens | Census Of Immune Cells | https://data.humancellatlas.org/explore/projects/cc95ff89-2e68-4a08-a234-480eca21ce79 |
| study15 | Mus musculus | Mapping the Mouse Cell Atlas by Microwell-Seq. | PMID:29474909 |
| study16 | Homo sapiens | Transcriptome Landscape of Human Folliculogenesis Reveals Oocyte and Granulosa Cell Interactions. | PMID: 30472193 |
| study17 | Homo sapiens | A Cellular Anatomy of the Normal Adult Human Prostate and Prostatic Urethra. | PMID: 30566875 |
| study18 | Homo sapiens | Single-cell reconstruction of the early maternalâ€“fetal interface in humans | PMID: 30429548 |
| study19 | Homo sapiens | Single-cell transcriptome analysis reveals differential nutrient absorption functions in human intestine | PMID: 31753849 |
| study20 | Homo sapiens | Single-Cell Transcriptomic Analysis of Primary and Metastatic Tumor Ecosystems in Head and Neck Cancer | PMID: 29198524 |
| study21 | Homo sapiens | Single-cell transcriptomic atlas of the human retina identifies cell types associated with age-related macular degeneration | PMID: 31653841 |

| | | | |
|---|---|---|---|
| study22 | Homo sapiens | Single-cell reconstruction of the adult human heart during heart failure and recovery reveals the cellular landscape underlying cardiac function | PMID:31915373 |
| study23 | Mus musculus | Single cell analysis reveals immune cell-adipocyte crosstalk regulating the transcription of thermogenic adipocytes | PMID: 31644425 |
| study24 | Mus musculus | An atlas of the aging lung mapped by single cell transcriptomics and deep tissue proteomics | PMID: 30814501 |
| study25 | Homo sapiens | The adult human testis transcriptional cell atlas | PMID: 30315278 |
| study26 | Homo sapiens | Single-cell reconstruction of follicular remodeling in the human adult ovary | PMID: 31320652 |
| study27 | Homo sapiens | Single-cell analysis of olfactory neurogenesis and differentiation in adult humans | PMID: 32066986 |
| study28 | Homo sapiens | Single-Cell Transcriptomic Map of the Human and Mouse Bladders | PMID: 31462402 |
| study29 | Mus musculus | Single cell analysis reveals immune cell-adipocyte crosstalk regulating the transcription of thermogenic adipocytes | PMID: 31644425 |
| study30 | Homo sapiens | Single-cell analysis of human adipose tissue identifies depot- and disease-specific cell types | PMID: 32066997 |
| study31 | Homo sapiens | Adipose tissue - Construction of a human cell landscape at single-cell level | https://www.nature.com/articles/s41586-020-2157-4 |
| study32 | Homo sapiens | Adrenal gland - Construction of a human cell landscape at single-cell level | https://www.nature.com/articles/s41586-020-2157-4 |
| study33 | Homo sapiens | Artery - Construction of a human cell landscape at single-cell level | https://www.nature.com/articles/s41586-020-2157-4 |
| study34 | Homo sapiens | Ascending colon - Construction of a human cell landscape at single-cell level | https://www.nature.com/articles/s41586-020-2157-4 |
| study35 | Homo sapiens | Bladder - Construction of a human cell landscape at single-cell level | https://www.nature.com/articles/s41586-020-2157-4 |
| study36 | Homo sapiens | Bone marrow - Construction of a human cell landscape at single-cell level | https://www.nature.com/articles/s41586-020-2157-4 |
| study37 | Homo sapiens | Cerebellum - Construction of a human cell landscape at single-cell level | https://www.nature.com/articles/s41586-020-2157-4 |
| study38 | Homo sapiens | Cervix - Construction of a human cell landscape at single-cell level | https://www.nature.com/articles/s41586-020-2157-4 |
| study39 | Homo sapiens | Small intestine duodenum - Construction of a human cell landscape at single-cell level | https://www.nature.com/articles/s41586-020-2157-4 |
| study40 | Homo sapiens | Appendix - Construction of a human cell landscape at single-cell level | https://www.nature.com/articles/s41586-020-2157-4 |
| study41 | Homo sapiens | Esophagus - Construction of a human cell landscape at single-cell level | https://www.nature.com/articles/s41586-020-2157-4 |
| study42 | Homo sapiens | Fallopian tube - Construction of a human cell landscape at single-cell level | https://www.nature.com/articles/s41586-020-2157-4 |
| study43 | Homo sapiens | Gallbladder - Construction of a human cell landscape at single-cell level | https://www.nature.com/articles/s41586-020-2157-4 |
| study44 | Homo sapiens | Heart - Construction of a human cell landscape at single-cell level | https://www.nature.com/articles/s41586-020-2157-4 |

### Supplementary Material

| | | | |
|---|---|---|---|
| study45 | Homo sapiens | Small intestine ileum - Construction of a human cell landscape at single-cell level | https://www.nature.com/articles/s41586-020-2157-4 |
| study46 | Homo sapiens | Small intestine jejunum - Construction of a human cell landscape at single-cell level | https://www.nature.com/articles/s41586-020-2157-4 |
| study47 | Homo sapiens | Kidney - Construction of a human cell landscape at single-cell level | https://www.nature.com/articles/s41586-020-2157-4 |
| study48 | Homo sapiens | Liver - Construction of a human cell landscape at single-cell level | https://www.nature.com/articles/s41586-020-2157-4 |
| study49 | Homo sapiens | Lung - Construction of a human cell landscape at single-cell level | https://www.nature.com/articles/s41586-020-2157-4 |
| study50 | Homo sapiens | Muscle - Construction of a human cell landscape at single-cell level | https://www.nature.com/articles/s41586-020-2157-4 |
| study51 | Homo sapiens | Omental adipose tissue - Construction of a human cell landscape at single-cell level | https://www.nature.com/articles/s41586-020-2157-4 |
| study52 | Homo sapiens | Pancreas - Construction of a human cell landscape at single-cell level | https://www.nature.com/articles/s41586-020-2157-4 |
| study53 | Homo sapiens | Peripheral blood - Construction of a human cell landscape at single-cell level | https://www.nature.com/articles/s41586-020-2157-4 |
| study54 | Homo sapiens | Lung pleura - Construction of a human cell landscape at single-cell level | https://www.nature.com/articles/s41586-020-2157-4 |
| study55 | Homo sapiens | Prostate - Construction of a human cell landscape at single-cell level | https://www.nature.com/articles/s41586-020-2157-4 |
| study56 | Homo sapiens | Rectum - Construction of a human cell landscape at single-cell level | https://www.nature.com/articles/s41586-020-2157-4 |
| study57 | Homo sapiens | Sigmoid colon - Construction of a human cell landscape at single-cell level | https://www.nature.com/articles/s41586-020-2157-4 |
| study58 | Homo sapiens | Spleen - Construction of a human cell landscape at single-cell level | https://www.nature.com/articles/s41586-020-2157-4 |
| study59 | Homo sapiens | Stomach - Construction of a human cell landscape at single-cell level | https://www.nature.com/articles/s41586-020-2157-4 |
| study60 | Homo sapiens | Brain temporal lobe - Construction of a human cell landscape at single-cell level | https://www.nature.com/articles/s41586-020-2157-4 |
| study61 | Homo sapiens | Thyroid - Construction of a human cell landscape at single-cell level | https://www.nature.com/articles/s41586-020-2157-4 |
| study62 | Homo sapiens | Trachea - Construction of a human cell landscape at single-cell level | https://www.nature.com/articles/s41586-020-2157-4 |
| study63 | Homo sapiens | Transverse colon - Construction of a human cell landscape at single-cell level | https://www.nature.com/articles/s41586-020-2157-4 |
| study64 | Homo sapiens | Ureter - Construction of a human cell landscape at single-cell level | https://www.nature.com/articles/s41586-020-2157-4 |
| study65 | Homo sapiens | Uterus - Construction of a human cell landscape at single-cell level | https://www.nature.com/articles/s41586-020-2157-4 |
